# QuickSeg: A fast, versatile and accurate algorithm for genomic copy number segmentation using dynamic programming

**DOI:** 10.64898/2026.08.31.747916

**Authors:** Balthasar Schlotmann, Francesco Favero, Alessio Locallo, Joachim Weischenfeldt

## Abstract

Copy number alterations are among the most common genomic aberrations in cancer and their accurate identification relies on robust segmentation of sequencing read-depth signals. Existing segmentation methods typically balance computational efficiency against segmentation accuracy and remain sensitive to technical artifacts present in sequencing data. Here, we present QuickSeg, a fast and versatile methodology that uses an exact dynamic programming algorithm to detect copy number segments using median-based error function. Motivated by the observation that sequencing depth distributions contain a small but pervasive population of outlying observations, this approach provides increased robustness to technical noise while simultaneously reducing the computational complexity of the segmentation problem. Across whole-genome sequencing of cancer cohorts, using breakpoint-supported somatic copy number alterations, we demonstrate improved segmentation precision over two widely used baseline methods, Circular Binary Segmentation (CBS) and Piecewise Constant Fitting (PCF), across a broad range of sensitivity thresholds. QuickSeg also consistently outperformed both methods with respect to runtime and memory usage. Collectively, our results show that robust median-based optimization provides both biological and computational advantages for copy number segmentation, enabling accurate analysis of large sequencing cohorts with minimal computational requirements.

## Background

Copy number alterations (CNAs) are a common form of structural variation in cancer genomes and frequently encompass large genomic regions containing hundreds to thousands of genes. Such alterations can drive tumour development through the loss of tumour suppressor genes or amplification of oncogenes ^1^. In contrast to single nucleotide variants, CNAs often have broad effects on gene dosage owing to their size and can therefore influence numerous biological pathways simultaneously. Many cancers are characterized by copy number instability (CIN), which results in recurrent gains and losses of whole chromosomes or chromosomal regions ^2^. Consequently, accurate detection of CNAs remains a fundamental task in cancer genome analysis.

Copy number segmentation is a special case of the broader changepoint detection problem, in which a sequential signal is partitioned into regions with distinct underlying properties ^3^. Numerous methods have been developed for this task, differing primarily in their assumptions about the data and in the trade-off between computational efficiency and segmentation accuracy. One of the most widely used approaches is Circular Binary Segmentation (CBS), which treats a chromosome’s copy number signal as a sequence of data points and searches for change-points that split it into segments of constant mean copy number ^4^.

In addition to heuristic approaches such as CBS, several methods aim to identify an optimal segmentation with respect to a predefined cost function, instead of recursively testing all breakpoint pairs. Dynamic programming algorithms provide an exact solution to this optimization problem but are often computationally expensive for large datasets. Approaches such as Piecewise Constant Fitting (PCF) (and related methods such as Pruned Exact Linear Time, PELT) treat the copy number signal as a sequence to be approximated by a step-wise function, using a piecewise-constant fit that minimises an error function ^5 6^. Despite these advances, copy number segmentation of large sequencing datasets remains computationally demanding and is often developed with one type of copy number data in mind (typically whole genome sequencing or arrays), and there is still a need for methods that combine robustness, accuracy, and computational efficiency.

Despite substantial methodological advances, accurate copy number segmentation remains challenging because sequencing data contain both biological signal and technical artifacts. Moreover, sparse copy number data from *e.g.* whole exome or panel sequencing poses a particular challenge to segmentation algorithms, and existing approaches often sacrifice computational efficiency for accuracy, or rely on heuristics that may affect segmentation quality. Here, we present QuickSeg, a fast and robust copy number segmentation algorithm that combines an *L*_1_ -loss framework with an exact dynamic programming solution. By leveraging the discrete structure of copy number states, QuickSeg achieves improved computational performance while maintaining optimal segmentation accuracy.

## Results

### Algorithm design

### Read-depth characteristics motivate an absolute-error formulation

Many copy number segmentation methods can be formulated as the minimization of a segmentation cost consisting of a fit term and a penalty for introducing additional breakpoints. Following the generalized formulation of the dynamic programming problem Killick et al. formulate in the PELT paper ^6^, a change-point analysis with *m* change points can be expressed as

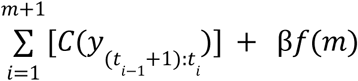

where

*y*_1_,…,*y_n_* are the observed data points, *t_i_* denotes the position of the ith breakpoint, C is the cost associated with an individual segment, and *f*(*m*) is a penalty term that discourages over-segmentation.

The choice of segment error function strongly influences both the statistical properties and computational characteristics of a segmentation algorithm. Many existing approaches use a squared-error (*L*_2_) loss ^5 6^, for which the optimal segment value is equal to the mean of the observations within the segment. This formulation is well suited to data that closely follows a Gaussian error model, but it is sensitive to outlying observations because large deviations are penalized disproportionately. Sequencing data, however, often contains technical artifacts that generate outlying observations. To reduce the influence of such artifacts, QuickSeg uses an absolute-error (*L*_1_) loss. A key property of the *L*_1_ loss is that the optimal segment value corresponds to the segment median rather than its mean. For symmetric distributions, the mean and median coincide, so the *L*_1_ – and *L*_2_ -loss are aligned in expectation, although they remain statistically distinct.

We therefore define the QuickSeg error function as

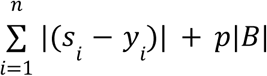

where *y_i_* is the observed read depth at position i, *s_i_* is the fitted segment value, *p* is the breakpoint penalty, and |*B*| denotes the number of breakpoints (or equivalently, the number of segments minus one).

### L1 segmentation enables an efficient dynamic programming formulation

A useful property of the absolute-error loss is that the optimal value for a segment corresponds to the segment median ^7^. Since the median must be one of the observed values within the segment, the set of possible segment values is finite. This observation forms the basis of the QuickSeg dynamic programming algorithm.

QuickSeg first determines the set of unique values observed in the input data, denoted *V*, which constitute the possible segment values. For each genomic position and each candidate value in *V*, the algorithm evaluates two alternatives: continuing the current segment or introducing a breakpoint and starting a new segment. A breakpoint is introduced whenever the cost of switching to a new segment, including the breakpoint penalty *p*, is lower than the cost of continuing the current segment.

During the forward pass, QuickSeg maintains the optimal score for each candidate segment value and records only the information required to reconstruct the optimal segmentation. Once all positions have been processed, the globally optimal solution is obtained by selecting the lowest-scoring final state and backtracking through the recorded breakpoint decisions. This procedure yields the exact optimum of the objective function.

Because the dynamic programming state space is restricted to the limited set of candidate segment values, runtime scales linearly with both the number of observations and the number of candidate states. After construction of the candidate value set, the computational complexity is *O*(*n*|*V*|). In the general case, identifying and sorting unique values requires *O*(*n log n*) preprocessing time, resulting in an overall complexity of *O*(*max*(*n*|*V*|, *n log n*)).

To reduce the preprocessing overhead associated with identifying unique values, the library implementations support user-defined candidate segment values. By explicitly defining the candidate state space, users can avoid the sorting step entirely while retaining the *O*(*n*|*V*|) segmentation runtime. The standalone implementation further provides an automatic state-space construction procedure based on a histogram of the expected read-depth range. For the vast majority of sequencing samples, this approach identifies all candidate states in linear time and therefore also avoids the cost of sorting.

In practice, sequencing depth data is characterized by a limited number of biologically meaningful read-depth levels, and the number of candidate states remains small relative to the number of observations. The theoretical worst-case complexity is encountered only when the number of unique values approaches the number of observations. This situation may arise following normalization procedures that transform integer read counts into high-precision floating-point values. For example, normalization against a matched control sample can introduce small numerical differences between observations that are biologically indistinguishable, substantially increasing the number of unique values without adding meaningful information to the segmentation problem. In such cases, the effective size of the state space can be reduced either by rounding observations to an appropriate precision or by providing a predefined set of candidate segment values. The former approach reduces the number of distinct states while preserving the exact optimization procedure, whereas the latter constrains the solution to a biologically meaningful set of segment values, but will not necessarily provide the optimal segmentation. Depending on the application, the latter can, however, act as a regularization mechanism by preventing segmentation between nearly identical copy-number states and thereby reducing sensitivity to numerical noise.

A common limitation of dynamic programming methods is their memory consumption, which often arises from storing intermediate scores and traceback information for all states. QuickSeg avoids storing the full dynamic programming score matrix and instead retains only the current score vector during the forward pass. For traceback, the algorithm stores the index of the globally optimal state at each position together with a single binary decision for each candidate state indicating whether a breakpoint was introduced. These decisions are represented as a compact bit vector, reducing memory requirements to *O*(*n*) integers and *O*(*n*|*V*|) bits while still permitting exact reconstruction of the optimal segmentation. As a result, QuickSeg achieves substantially lower memory usage than conventional dynamic programming implementations despite computing the exact optimum of the segmentation objective. We also implemented additional optimizations on AVX512 capable CPUs described in a supplemental note.

### Benchmark comparison to existing segmentation algorithms

Numerous software packages for copy number analysis have been developed, often integrating normalization, bias correction, segmentation, visualization, and downstream interpretation into a single workflow. While such integrated frameworks are valuable for routine analysis, they complicate the evaluation of individual segmentation methods because differences in performance may arise from upstream or downstream processing steps rather than the segmentation algorithm itself. The distinction between a segmentation algorithm and a complete copy number analysis workflow is not always maintained in the literature. As a result, benchmarking studies may inadvertently compare entire analysis pipelines rather than the underlying segmentation methods. Because the objective of the present study was to evaluate segmentation performance specifically, we restricted comparisons to standalone segmentation algorithms that can be assessed independently of normalization, bias correction, and downstream copy-number analysis steps. This substantially limited the pool of eligible methods, a challenge also noted by Zhang et al., who reported that several commonly used segmentation approaches, such as HMM, are available only as components of larger analysis frameworks ^8^. We therefore selected CBS and PCF, implemented by the R packages “DNAcopy” and “copynumber”, respectively ^9 5^. These are two widely used segmentation algorithms that are available as standalone implementations and can be evaluated independently of preprocessing and downstream analysis steps.

### Simulated data shows clear runtime and memory usage improvements

We first evaluated performance using simulated data generated from piecewise-constant signals with stochastic sampling noise. All evaluated methods recovered the simulated breakpoint structure using their default parameters. However, we deliberately did not attempt to construct a challenging synthetic accuracy benchmark, as such datasets are inherently shaped by the assumptions of the simulation model and may therefore favour particular classes of segmentation algorithms. Instead, simulated data were used primarily to evaluate computational performance, since runtime and memory consumption can be assessed objectively while controlling the number of observations and candidate segment values. The simulation datasets were composed from 100 to 1,000,000 points. Because the runtime of QuickSeg depends on the number of candidate segment values, benchmarks were performed using an unconstrained state space in which candidate values were obtained by sorting and extracting the distinct observed values from the data and a constrained state spaces containing 64 candidate states to evaluate the impact of the CPU optimization (AVX-512, see methods), or 63 for the closest comparison without the optimizations.

Our results demonstrate that QuickSeg consistently outperformed both CBS and PCF with respect to runtime (Fig. 1A) and memory (Fig. 1B) consumption across all simulated dataset sizes. CBS was the slowest method throughout the benchmark, while PCF achieved faster runtimes but remained considerably slower than QuickSeg.

**Figure 1.**
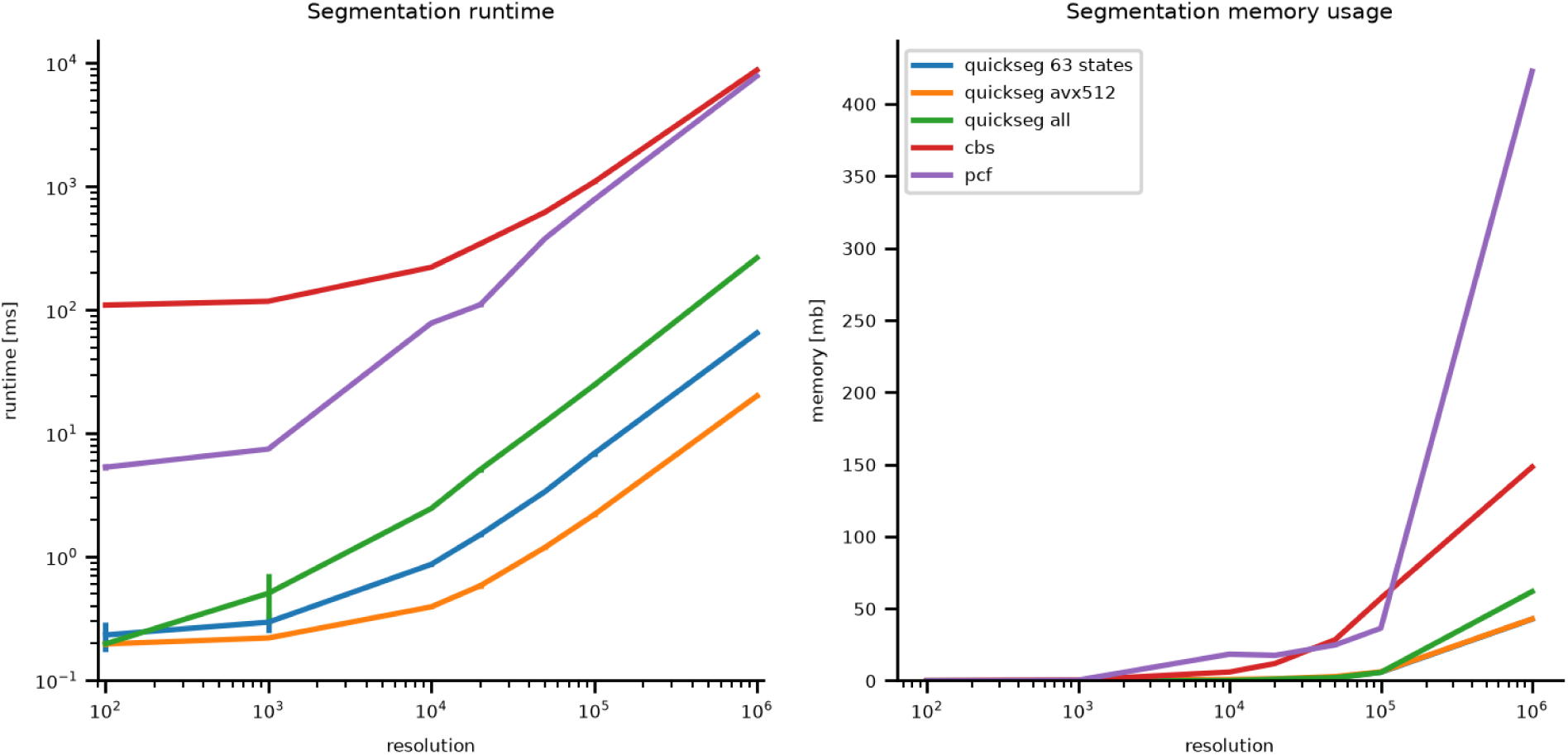
Runtime and memory usage of QuickSeg, PCF and CBS. A, Runtime of CBS, PCF, and QuickSeg across simulated datasets of increasing size. B, Peak memory consumption of CBS, PCF, and QuickSeg across the same datasets.

Across all simulated datasets, QuickSeg required only approximately 2% and 4% of the runtime of CBS and PCF, respectively. The AVX-512 optimized implementation using 64 candidate states was consistently faster than the otherwise equivalent 63-state version, reaching only 40% of the runtime at one million datapoints. As expected, runtime scaled approximately linearly with the number of candidate segment values.

Memory consumption showed a similar pattern. At a resolution of one million datapoints, QuickSeg required only 137Mb memory, which corresponded to only 60% and 20% of the memory used by CBS (218Mb) and PCF (666Mb). This reduced memory footprint reflects the compact dynamic-programming representation employed by QuickSeg and remained substantially lower than that of the competing methods across all benchmark sizes.

### QuickSeg shows more precise segmentation on the TCGA prostate cancer cohort than CBS and PCF

We evaluated segmentation performance using the TCGA prostate cancer cohort. Prostate cancer genomes frequently exhibit extensive CNAs and structural variation (SV), making them a suitable model system for benchmarking copy number segmentation methods. Pre-binned read-depth data and previously identified SV calls were used for all analyses ^10^.

A major challenge in evaluating copy number segmentation algorithms is the absence of a universally accepted ground truth. Previous studies have used Multiplex Ligation-dependent Probe Amplification (MLPA) measurements for validation ^11 5^. However, because MLPA interrogates only a predefined set of genomic loci, it provides only partial information regarding genome-wide segmentation performance ^12^. Furthermore, to our knowledge, the extent to which MLPA measurements represent a superior ground truth relative to modern whole-genome sequencing-based copy number analyses has not been systematically evaluated. More generally, establishing an objective benchmark for copy number segmentation remains difficult because the true underlying segmentation of cancer genomes is rarely known.

To address this limitation, we evaluated segmentation performance using independently identified SV breakpoints. While balanced SVs such as reciprocal translocations and inversions do not contribute to copy number breakpoints and SVs themselves can be subject to false positive and false negative calls, they provide an orthogonal source of support that is independent of read-depth based segmentation. Consequently, agreement between predicted copy number breakpoints and SV breakpoints can be used as an objective measure of segmentation performance without inherently favouring any of the evaluated algorithms.

Because not all copy number changes are associated with detectable SVs, and because SV callsets are themselves imperfect, absolute performance estimates are likely conservative. Nevertheless, this approach provides a practical and biologically meaningful framework for comparing segmentation algorithms on real tumour genomes. We, therefore, used this approach to evaluate the computational performance on the TCGA prostate cancer cohort (Fig. 2). Consistent with the results obtained on simulated data, QuickSeg was the fastest method across all samples (Fig. 2B). Runtime remained below 10 seconds in the worst case. The performance of PCF proved substantially more sensitive to parameter choice than that of CBS. While favourable parameter settings allowed PCF to outperform CBS, its runtime varied considerably and worst-case performance was often substantially slower.

**Fig. 2.**
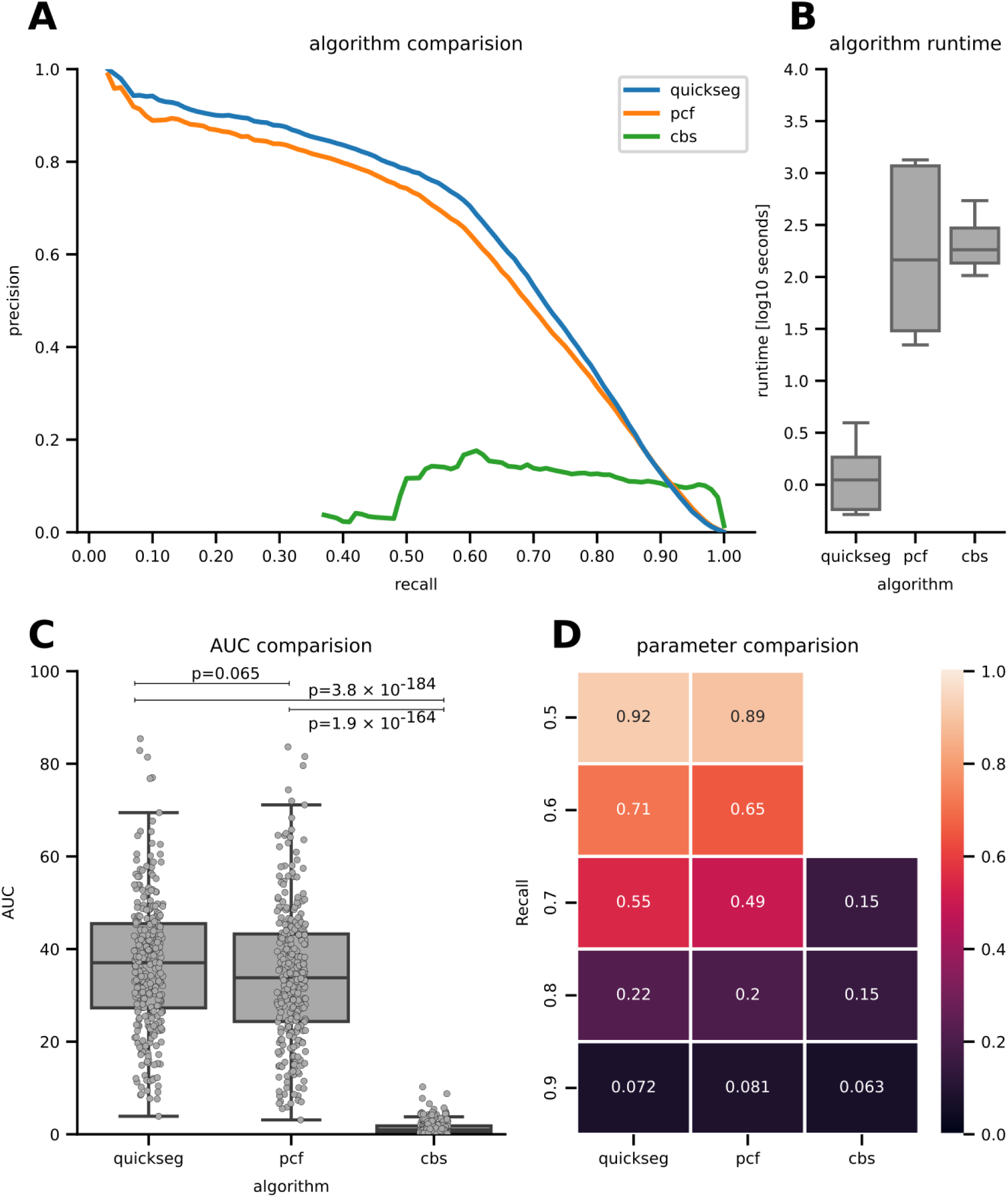
Benchmarking on TCGA prostate cancer whole-genome sequencing data. A) Precision-recall curves for QuickSeg (blue), PCF (orange), and CBS (green) generated by varying the segmentation penalty. Agreement between segmentation breakpoints and independently identified SV breakpoints was used as the performance metric. B) Runtime performance across the TCGA prostate cancer cohort. Boxes indicate the interquartile range, center lines the median, and whiskers extend to the farthest datapoint within 1.5 times the interquartile range. C) Distribution of sample-level (N=302) area under the precision-recall curve values for each algorithm. Boxes indicate the interquartile range, center lines the median, and whiskers extend to the farthest datapoint within 1.5 times the interquartile range. D) Heatmap showing the precision values obtained for parameter settings producing approximately matched median sensitivities of 50%, 60%, 70%, 80%, and 90% across samples. CBS did not achieve median sensitivities of 50% or 60% and was therefore excluded from these comparisons.

**Fig 3.**
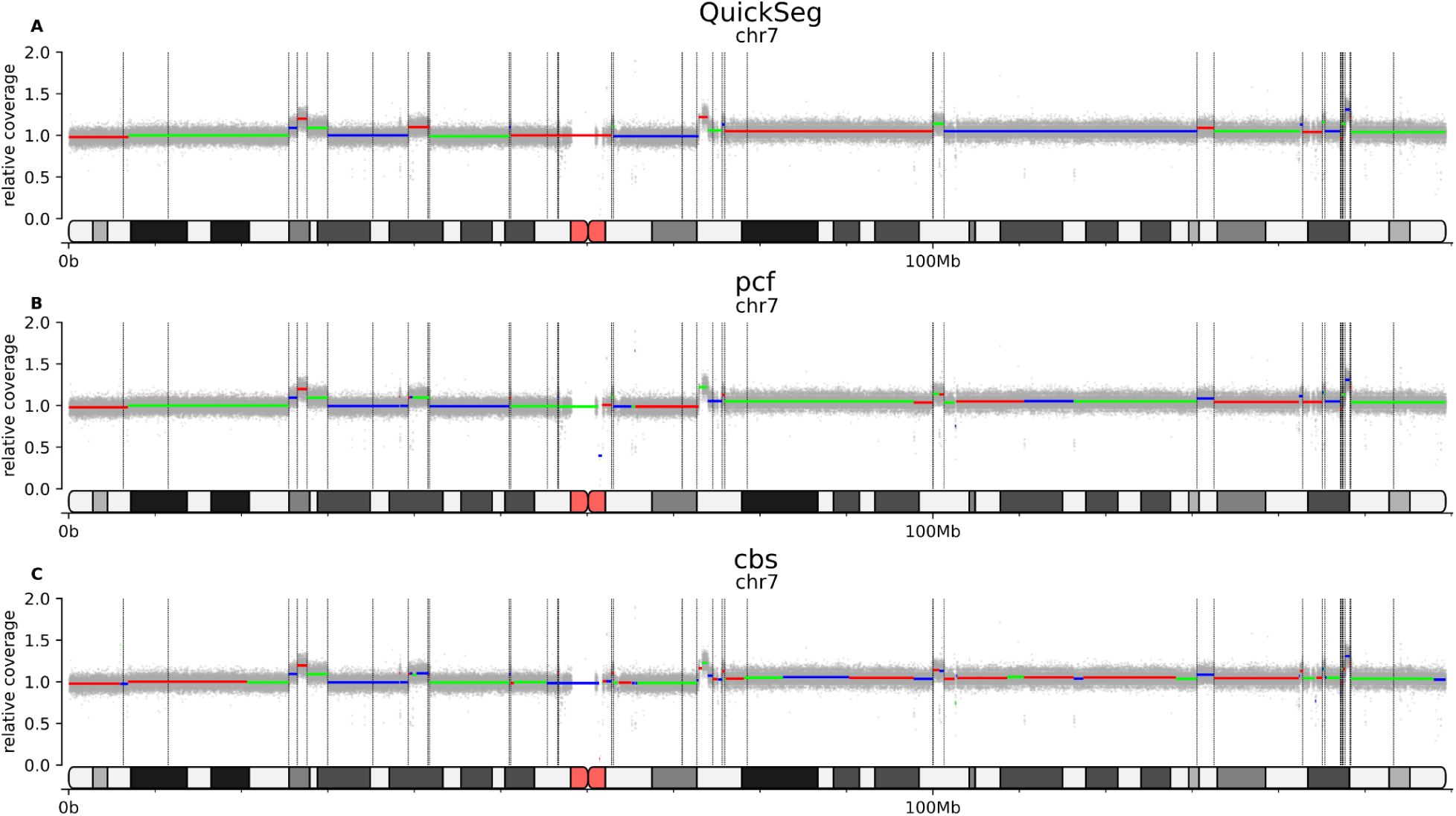
Matched sensitivity show QuickSeg has improved precision over pcf and cbs. A, Segmentation using Quickseg. B, Segmentation using PCF. C, Segmentation using CBS

**Fig 4.**
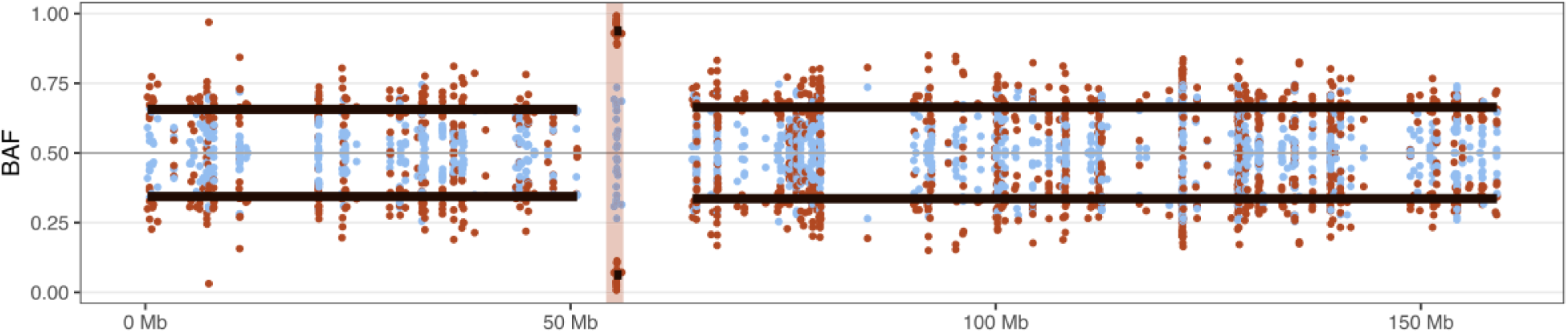
Quickseg shows applicability across different input data. A, Quickseg being used to segment allele frequency (black lines) based on scRNA-seq data of chromosome 7 from glioblastoma. B-allele frequency (BAF) is shown on the y-axis and genomic positions of chromosome 7 on the x-axis. Blue dots are from a normal-comparison sample and red dots are from the glioblastoma sample from the same patient.

For each algorithm, we generated precision-recall curves by varying the segmentation penalty. We deliberately avoided receiver operating characteristic (ROC) curves, as copy number segmentation is an extremely imbalanced classification problem in which every genomic bin constitutes a potential breakpoint while only a small fraction corresponds to true events. Under these conditions, specificity is dominated by the overwhelming number of true negative observations and may therefore provide an overly optimistic assessment of performance. Precision-recall curves provide a more informative summary of breakpoint detection accuracy and have similarly been adopted in previous CNV benchmarking studies such as Gabrielaite et al. ^11^.

Across the evaluated parameter ranges, QuickSeg achieved consistently higher performance than both CBS and PCF (Fig. 2A) and achieved the highest area under the precision-recall curve (AUC) (Fig. 2C). CBS exhibited comparatively limited responsiveness to parameter changes and showed a tendency toward over-segmentation, resulting in a restricted range of attainable precision and sensitivity values and a low AUC. In contrast, both PCF and QuickSeg spanned a much broader range of operating points.

As expected, performance differences between algorithms diminished at very low and very high sensitivities, corresponding to the extreme ends of the precision-sensitivity curve. Across the practically relevant range of approximately 10% to 90% sensitivity, however, QuickSeg consistently achieved higher mean precision than PCF (min = -0.1%pt., max = 6%pt.). Consequently, QuickSeg delivered superior performance throughout most of the evaluated precision-sensitivity space, indicating a more favourable trade-off between breakpoint detection and over-segmentation.

To facilitate direct comparison between algorithms, we additionally identified parameter settings yielding approximately matched median sensitivities of 50%, 60%, 70%, 80%, and 90% across all samples (Fig. 2D). For each sensitivity level, precision was then compared between segmentation methods. This analysis confirmed the trend observed in the precision-sensitivity curves, with QuickSeg achieving higher precision than both CBS and PCF across most operating points. QuickSeg outperformed the competing methods at all evaluated sensitivities up to 90%. At the highest sensitivity level, PCF achieved marginally higher precision, with an advantage of approximately one percentage point. However, precision was below 10% for all algorithms at this operating point, suggesting limited practical utility of such highly sensitive parameterisations.

CBS exhibited comparatively limited parameter responsiveness and no parameter setting produced median sensitivities near 50% or 60%. Consequently, CBS was excluded from these comparisons. Even at higher sensitivity levels, CBS generally achieved lower precision than either QuickSeg or PCF, consistent with its tendency towards over-segmentation observed throughout the benchmark.

### Applications

The QuickSeg R library has been used by Hendriksen et al. in the development of ecsingle, a tool for the detection of copy number alterations and ecDNA from 10X single cell RNA-seq data ^13^. In this manuscript, we used QuickSeg to segment allele frequencies obtained from brain cancer-derived 10X single cell RNA-sequencing reads to identify regions with likely ecDNA, demonstrating QuickSeg’s versatility beyond read depth segmentation.

A variant of the QuickSeg method has recently been implemented in the sequenza R package (https://bitbucket.org/sequenzatools/sequenza). This implementation retained the core algorithm for scoring penalties, but extended it to operate on changes in a multi-dimensional array rather than a single vector of data. This approach allows detection of allele-specific copy-number changes, when coverage/depth ratios and corresponding B-allele frequencies are evaluated in the array, as well as multi-sample copy-number changes, when multiple samples from the same donor are evaluated in the array.

## Discussion

In this study, we presented QuickSeg, a copy number segmentation algorithm based on exact dynamic programming and an absolute-error loss formulation. Across both simulated and real sequencing data, QuickSeg consistently demonstrated substantially lower runtime and memory requirements than CBS and PCF while achieving improved segmentation performance. These results indicate that robust and computationally efficient copy number segmentation can be achieved without relying on breakpoint heuristics or approximate optimization procedures.

The primary limitation of the present benchmark is the absence of a universally accepted ground truth for copy number segmentation. Although independently identified SV breakpoints provide a substantially more objective validation framework than approaches based on targeted assays or manually curated examples, the correspondence between SVs and copy number alteration is inherently incomplete. Some SVs are copy number neutral and therefore should not be expected to generate detectable copy number breakpoints, while conversely many copy number changes may arise without an identifiable SV call. In addition, current SV callsets contain both false positive and false negative events, further limiting the achievable agreement between segmentation and SVs.

These limitations imply that the absolute performance of all evaluated segmentation methods is likely underestimated. Indeed, visual inspection frequently revealed clear copy number transitions without accompanying structural variant calls as well as structural variant breakpoints occurring in otherwise uniform copy number regions. However, because SV detection is independent of read-depth based segmentation, these imperfections are not expected to systematically favour any particular algorithm. We therefore consider agreement with SV breakpoints to provide a conservative but comparatively unbiased measure of segmentation performance.

Conversely, the same principle may be applied in the opposite direction, using robust copy number segmentation as an orthogonal source of evidence when evaluating SV calling methods. The development of benchmarking frameworks based on multiple independent genomic signals may therefore provide a more reliable alternative to the single-modality benchmarks that have historically dominated the field. We demonstrate the versatility of QuickSeg, by applying and robustly detecting both large and smaller CNAs from extremely sparse data such as scRNA-seq data, which depends on genotyped SNPs present in the 10X scRNA-seq data. Future work will be required to compare this versatility with existing algorithms such as CBS and PFS.

From a computational perspective, QuickSeg occupies an intermediate position between classical change-point dynamic programming algorithms and finite-state decoding methods such as the Viterbi algorithm ^14 15^. Like Viterbi decoding, the algorithm performs optimization over a finite set of states and updates these states sequentially along the genome. However, unlike hidden Markov models, the state space is not fixed in advance and no explicit probabilistic transition model is required, since all state transitions are treated uniformly through a single breakpoint penalty. Instead, the candidate states emerge naturally from the median property of the absolute-error loss itself. Conversely, while QuickSeg targets the same optimal piecewise-constant segmentation sought by classical dynamic programming approaches, it reduces the search space by exploiting the limited number of biologically meaningful copy-number states rather than by pruning candidate breakpoint locations. To our knowledge, this formulation has not previously been exploited in copy-number segmentation and offers an alternative route to computational acceleration that is complementary to existing strategies.

As a result, QuickSeg combines the computational efficiency of finite-state algorithms with the exact optimization framework of penalized segmentation, avoiding both the quadratic breakpoint search of classical segmentation algorithms and the quadratic state-transition step of general finite-state decoders and suggests an alternative perspective on change-point detection in which optimization is performed over candidate states rather than breakpoint locations.

## Data availability

The TCGA prostate cancer data used in this study are available through The Cancer Genome Atlas (TCGA) data portal https://portal.gdc.cancer.gov/. Source code for the QuickSeg segmentation algorithm is available as open-source software through dedicated R and Python libraries as well as a standalone Rust implementation at:

R package: https://github.com/Balthasar-eu/Rquickseg

Python package: https://github.com/Balthasar-eu/pyquickseg

Rust standalone implementation: https://github.com/Balthasar-eu/quickseg

Benchmarking scripts: https://github.com/Balthasar-eu/quickseg_benchmarking

## Methods

### Synthetic runtime performance benchmark

For synthetic benchmarks, copy-number profiles were generated as piecewise-constant signals consisting of five segments. Observations within each segment were sampled independently from binomial distributions with parameters Bin(200, 0.5), Bin(300, 0.5), Bin(200, 0.5), Bin(100, 0.5), and Bin(200, 0.5), resulting in segment means of approximately 100, 150, 100, 50, and 100 reads, respectively. Segment lengths were fixed at 40%, 10%, 20%, 15%, and 15% of the total profile length. Datasets containing 100, 1,000, 10,000, 20,000, 50,000, 100,000, and 1,000,000 observations were generated to evaluate scaling behaviour across a wide range of problem sizes.

Runtime benchmarking was performed using standalone scripts for each segmentation algorithm executed on identical input datasets. To minimize the influence of implementation-specific file I/O overhead, runtime measurements were started after data loading and terminated immediately after segmentation. Consequently, the reported runtimes reflect only the execution time of the segmentation algorithms themselves and do not include file parsing or output generation. Each benchmark was repeated five times and the median runtime was used for subsequent analysis. Memory consumption was measured using GNU time (-f %M), which reports the peak resident memory usage of the process. To avoid counting memory used by the interpreter and dependencies, we subtracted the memory usage for 100 datapoints from all measurements, ensuring all algorithms have the same starting point. The Python implementations of QuickSeg using 63, 64, and unrestricted candidate segment states were included in the synthetic benchmarks to evaluate the effects of state-space size and AVX-512 optimization.

### TCGA data

For evaluation on real data, we analyzed the TCGA prostate cancer cohort comprising 407 tumour samples. SV calls were originally called using the Manta structural variant caller ^16^ and obtained directly from the TCGA repository.

Seventy-two samples originating from sequencing centre 02 were excluded because they consistently exhibited quality-control issues and contained unusually low numbers of structural variant (SV) calls. An additional 33 samples were removed because the limited number of SV breakpoints prevented meaningful assessment of segmentation performance. In these samples, even the best-performing segmentation result from any evaluated algorithm achieved less than 50% precision, indicating insufficient benchmark information rather than poor algorithmic performance. After filtering, 302 samples remained for downstream analysis. Read-depth data were analysed using pre-computed 10 kb genomic bins.

### Benchmarking

Segmentation performance was assessed by comparing predicted copy-number breakpoints with independently identified SV breakpoints. Two breakpoints were considered concordant if they occurred within 50 kb of one another, corresponding to five bins on either side of the predicted breakpoint. Duplicate SV calls occurring within the same bin were collapsed prior to analysis because they cannot be resolved at the chosen binning resolution.

To reduce the number of balanced structural variants and enrich for copy-number-associated events, a 20 kb window centred on each breakpoint was tested using a Kolmogorov-Smirnov test comparing the coverage distributions on either side of the breakpoint. An SV was retained if at least one of its two breakpoints exhibited a significant coverage change (p < 0.05). This procedure preferentially retains SVs associated with detectable copy-number alterations while removing many balanced rearrangements that are not expected to create copy-number breakpoints.

For each sample, breakpoints were classified as shared between the segmentation and SV callsets, unique to the SV callset, or unique to the segmentation callset. Shared breakpoints were treated as true positives (TP), while SV-only and segmentation-only breakpoints were treated as false negatives (FN) and false positives (FP), respectively. Precision was calculated as TP/(TP+FP) corresponding to the fraction of predicted breakpoints supported by an SV call, and sensitivity as TP/(TP+FN), corresponding to the fraction of SV breakpoints recovered by the segmentation. Chromosome boundaries were treated as shared breakpoints in all samples to ensure consistent behaviour at the extreme ends of the precision-sensitivity curves.

For each sample, overall segmentation performance was summarized as the mean interpolated precision across the evaluated sensitivity range.

Real-data analyses were performed using the standalone Rust implementation of QuickSeg using all unique values as states, while PCF and CBS were executed through the copynumber and DNAcopy R packages, respectively. QuickSeg and PCF were applied to linearized read-depth data, whereas CBS was run on log-transformed values, consistent with its recommended input format. To characterize performance across a wide range of operating points, each algorithm was evaluated over a broad range of penalty parameters. QuickSeg was run with penalties ranging from 0.01 to 10^6^, PCF with penalties ranging from 0.001 to 10^6^, and CBS with significance thresholds ranging from 0.1 to 10^−8^.

### AVX-512 optimization

For applications with 64 candidate segment values, QuickSeg provides a specialized implementation based on the AVX-512 instruction set. In this case, all candidate segment values and their associated dynamic-programming scores fit entirely within the processor’s 512-bit vector registers. As a result, the complete dynamic-programming state can be maintained in CPU registers throughout the forward pass.

Unlike conventional implementations, which repeatedly load and store intermediate score vectors in memory, the AVX-512 implementation requires only the newly observed data point to be loaded at each iteration. Candidate segment values remain resident in registers, while score updates, breakpoint decisions, absolute-difference calculations, and minimum-score reductions are performed using vectorized instructions. The absolute-error term is calculated through bitwise masking of the sign bit, avoiding additional arithmetic operations, and the global minimum score is determined using a binary search entirely within the registers containing the scores.

This optimization effectively eliminates memory traffic for the dynamic-programming state. On AVX-512 capable processors, this results in approximately twice as fast segmentation when using 64 candidate states compared to 63 candidate states, for which this optimization is not implemented. When combined with the user-defined candidate state mode, this implementation can substantially accelerate segmentation tasks.

## Author contributions

B.S. Conceived and implemented the algorithm. A.L. and F.F. provided development feedback and tested the implementation. F.F implemented the sequenza variant of the algorithm. B.S., F.F. and J.W. wrote the manuscript. J.W. and F.F. supervised the work.

## Acknowledgements

The results shown here are based upon data generated by the TCGA Research Network: https://www.cancer.gov/tcga.

